# Mapping Genome-wide Transcription Factor Binding Sites in two *Rhodanobacter* strains Isolated from Extreme Environments

**DOI:** 10.64898/2026.06.06.730518

**Authors:** Vishalakshi Bhanot, Alexey Kazakov, Guilherme M. V. de Siqueira, Alex Codik, Shweta Priya, Anne Kakouridis, Valentine V. Trotter, Adam M. Deutschbauer, Ian Blaby, Leo Baumgart, Aindrila Mukhopadhyay

## Abstract

Identification of target genes for bacterial transcription factors (TFs) provide a deeper understanding of bacterial response to environmental conditions. We focused on two *Rhodanobacter* sp. isolates from the subsurface of the U.S. Department of Energy Oak Ridge Reservation (ORR), an environment characterized by high-concentration mixed-waste contamination plumes. Bacterial strains in this subsurface ecosystem experience changing and complex environmental conditions, including variable nutrient and oxygen availability, fluctuating pH, nitrate and metals. We examined 91 DNA-binding TFs from 17 TF families in two *Rhodanobacter* strains, *R. denitrificans* FW104-10B01 and *R. thiooxydans* FW510-R12, using both DNA affinity purification sequencing (DAP-seq) and comparative genomics. Target genes regulated by 31 TFs were identified. Despite being isolated from different wells, the two strains shared a highly similar core regulatory network, with orthologous TFs regulating nearly identical sets of genes. These conserved networks governed essential survival functions, including nutrient uptake (e.g., phosphate), carbohydrate metabolism, motility, and stress responses to heavy metals and oxidative environment. TF-binding sequence motifs were also inferred for each of the 31 TFs. Key results include the elucidation of a global regulator Clp which regulates a large number of genes, particularly those involved in biofilm formation and motility. Functional validation through RT-qPCR confirmed that Clp activates the expression of genes critical for type IV pilin structure and flagellar regulation. Our work highlights the advantages of synergistically employing DAP-seq and comparative genomics, which enabled us to corroborate findings and obtain a larger view of the regulatory landscape of these non-model denitrifying bacteria.

## Introduction

Bacteria face numerous stressors in the complex and dynamic environments they inhabit. To survive in these constantly changing conditions, they rely on transcriptional regulation to mount survival responses, with transcription factors (TFs) playing a crucial role in orchestrating these processes. TF function depends on its ability to recognize specific DNA sequences or TF binding sites (TFBSs); short conserved DNA regions located in regulatory regions of target genes that facilitate or repress the expression of their respective target genes by either recruiting and stabilizing the RNA polymerase at the promoter region or preventing its binding, respectively, leading to differential expression of genes (1–3). Studying TFs enables the mapping of complex regulatory networks and the corresponding target genes which in turn provides insight into how bacteria adapt to dynamic and multifaceted environments (4). Bacterial signaling has been deeply studied for gut bacteria, and for bacteria isolated from different environments, including sites contaminated with highly toxic amounts of several stressors (5–7), and such investigations enabled better understanding of microbial dynamics in these environments (8).

The aquifer in the U.S. Department of Energy’s Oak Ridge Reservation (ORR), Tennessee represents a unique ecosystem characterized by varied gradients of pH, nitrate, metal radionuclides, heavy metals, and organic contaminants (9). Denitrifying bacteria from the genus *Rhodanobacter* are abundant and active in the groundwater and sediment subsurface of this environment where heavy metals and acidic conditions are key factors shaping the community structure (9,10). In 2012, the species *Rhodanobacter denitrificans* was first described after the isolation of strains 2APBS1^T^ and 116-2 from the ORR site (11), and the continued efforts to characterize *Rhodanobacter* isolates at the (pan) genomic level since then reflect the variability of features across strains and the importance of horizontal gene transfer events for the distribution of tolerance mechanisms in this environment (10,12). However, the specific regulatory networks involved in adaptation of *Rhodanobacter* isolates in the context of stressors present at the ORR remain largely underexplored.

In this study we used a combination of DNA affinity purification sequencing (DAP-seq) and comparative genomics to examine such regulatory features. DAP-seq is next-generation sequencing of fragmented genomic DNA libraries after enrichment using *in vitro* expressed TFs (13,14). TFs from two *Rhodanobacter* strains isolated from different environments at the ORR site were investigated in this study: *R. denitrificans* FW104-10B01, isolated from a well with elevated concentrations of stressors such as nitrate, sulfate, and uranium, and *R. thiooxydans* FW510-R12, which was isolated from a more acidic site where other stressors are not as abundant (10). Despite differences in their spatial distribution, these two strains are evolutionarily closely related. A previous analysis revealed that these two strains share greater than 89% average amino acid identity (AAI) and average nucleotide identity (ANI) (10), which can suggest similarities in their regulatory networks despite differences in their biological niches. In this study, we combined DAP-seq and comparative genomics approaches to obtain a deeper assessment of the regulatory networks of these two *Rhodanobacter* strains.

## Materials & Methods

### Bacterial strains and growth conditions

Two *Rhodanobacter* strains, *Rhodanobacter denitrificans* FW104-10B01 and *Rhodanobacter thiooxydans* FW510-R12, isolated and archived from contaminated groundwater wells in the U.S. Department of Energy Field Research Center in ORR, were used for this study. The details of their isolation and their overall genomic features have been reported previously (10). Transposon mutants for the global TF *clp* in *R. denitrificans* FW104-10B01 (tn::*clp*), were obtained from a previously constructed RB-TnSeq library (15–17).

The bacteria were recovered from frozen glycerol stocks by streaking onto Reasoner’s 2A medium (R2A) agar plates (HiMedia Laboratories LLC, Kennett Square, PA, USA) and incubated at 30 °C (9,18). Primary cultures were prepared by inoculating single colonies of *Rhodanobacter* strains into 1 mL of R2A broth and incubating at 30 °C with shaking at 200 rpm overnight. The following day, secondary cultures were established by inoculating the primary culture into fresh R2A medium at a 1:100 (v/v) dilution. Secondary cultures were incubated under the same conditions (30 °C, 200 rpm) and used as inocula for kinetic growth assays. In growth experiments, secondary cultures were adjusted to an initial optical density at 600 nm (OD_600_) of 0.1 in R2A medium and dispensed into 48-well flat-bottom microtiter plates (Corning Falcon) at a final volume of 300 μL per well. Plates were sealed with semi-permeable Breathe-Easy® membranes (Sigma-Aldrich, USA Scientific) and incubated at 30 °C in a SpectraMax M2e microplate reader (Molecular Devices, San Jose, CA) with continuous agitation. OD_600_ was recorded every 10 min for up to 60 h. For the transposon mutant tn::*clp* the growth medium was supplemented with kanamycin at a final concentration of 5 μg mL^-1^. In each case, growth measurements were performed for three biological replicates, each with three technical replicates. Growth data were processed and analysed using GraphPad Prism (Version 10.4.1).

### Genomic DNA Extraction, Library Preparation and TF Preparation for DAP-seq

A set of 91 TFs from *Rhodanobacter* strains were selected for DAP-seq experiments. The TF selection aimed to ensure broad functional relevance, including nutrient uptake, motility, biofilm formation, and stress response, as well as TF family diversity, encompassing both common and strain-specific TFs. Genomic DNA (gDNA) was extracted from freshly grown cultures of *R. denitrificans* FW104-10B01 and *R. thiooxydans* FW510-R12 using DNeasy PowerSoil Pro Kit (Qiagen, Redwood City, CA, USA), as per manufacturer’s instructions, and prepared via fragmentation to an average fragment size of 75 bp to generate libraries using the KAPA HyperPrep Kit followed by 10 cycles of PCR amplification, according to the protocol described in Baumgart *et al* 2021 (14).

A workflow at the Joint Genome Institute (JGI, Berkeley, CA) was employed for obtaining *in vitro* expressed TFs for the DAP-seq assays. In this workflow, TF protein sequences were reverse translated using the BOOST design software with an *Escherichia coli* codon frequency table (19). Specifically, the “balanced” strategy was used for codon refactoring, which aims to match the frequency of codons to that of the *E. coli* genome. DNA linkers (30mers) were added to the 5’ and 3’ of each DNA sequence enabling scarless, in-frame, assembly downstream of a HALO-tag into pIX-Halo_PaqCI, linearized by PaqCI digest. Linear synthetic DNA fragments (Twist BioScience, CA, USA) were PCR amplified and assembled into the linearized vector using the NEBuilder Hi-Fi Assembly kit (New England BioLabs, Ipswich, MA, USA). All constructs were verified by sequencing using the Pacific Biosciences Sequel IIe platform and analyzed using custom pipelines. Details of the sequences are provided in **Supplementary Table 1**.

### DAP-seq Assay, Raw Data Processing and Motif Discovery

DAP-seq experiments were carried out following the multiDAP method described in Baumgart *et al* 2021 (14), with minor modifications for Halo-tagged TFs. In brief, *in vitro* expressed Halo-tagged TFs from each *Rhodanobacter* strain were incubated with 10 μL Magne HaloTag Beads (Promega, Madison, WI, USA) and the multiplexed genomic DNA libraries of both species. The resulting DNA-TF complexes were then washed, recovered, amplified, and sequenced to identify the enriched binding sites in the genome.

Sequenced reads were adapter trimmed and quality filtered using BBTools version 38.96 (https://bbmap.org/) using the following options: k=21 mink=11 ktrim=r tbo tpe qtrim=r trimq=6 maq=10. Filtered reads were aligned to the reference genome of both *R. denitrificans* FW104-10B01 (RefSeq Accession No. NZ_CP088922.1) and *R. thiooxydans* FW510-R12 (RefSeq Accession No. NZ_CP088923.1) using bowtie2 (20) version 2.4.2 with the options --no-mixed --no-discordant. Peaks were called using MACS3 version 3.0.0a6 (21), using combined negative control samples with mock protein expression as the background file, and the following options: --call-summits --keep-dup 1, and --gsize with the total reference genome size. Sequences representing up to the 100 strongest peaks, as scored by signal value in column 7 of the narrowPeak files, were extracted using a custom python script. These peak sequences were used to generate motifs using MEME (MEME Suite v.5.3.0) (22) with a zero-order background file generated from the reference genome, and the following additional options: -dna -revcomp -mod anr -nmotifs 2 -minw 8 -maxw 32. Motif calling was run twice for each dataset, once with the entire peak regions and once with only the regions +/-30 bp flanking the summit location of each peak. Results of the peak-calling step were output in the BED file format for input into Bedtools 2.28 (23) for peak analysis in conjunction with the *Rhodanobacter* GFF files. Annotated peaks were then sorted by position using the sort-bed utility in BEDtools. Sorted peaks were input into the bedmap utility from the BEDOPS 2.4.39 program (24) to calculate statistics and overlaps for all the putative annotations.

### TF-binding Motif Validation with Comparative Analysis of Evolutionary Conservation

Comparative genome analysis was used if TF binding motif prediction by MEME from DAP-seq peak sequences did not give satisfactory results. The comparative genomics method assumes higher evolutionary conservation of true regulatory sites in comparison to non-functional sequences (25). Thus, if a DAP-seq peak with the highest FCE (fold change enrichment) value corresponds to a functional TF binding site, similar TF binding sites are expected to be found upstream of orthologous genes in closely related genomes, if these genomes encode orthologous TFs. Then, a conserved candidate TF-binding motif can be inferred from these orthologous upstream regions and used for scoring of DAP-seq peaks. For the comparative analysis, we downloaded 82 *Rhodanbacter* spp. genomes with annotated genes from GenBank (26) (listed in **Supplementary Table 2**). For each TF, we picked a DAP-seq peak with the highest read coverage in the non-coding region and identified the gene located immediately downstream from the peak in both directions. Orthologous proteins were identified with Orthofinder v. 2.5.5 (27). Non-coding upstream regions of orthologous genes were extracted from the genomes using Biopython (28). Upstream regions shorter than 50 bp were excluded from the analysis, because they may represent intergenic spacers within operons. If, after that, a set of orthologous upstream regions contained less than 20 sequences, the gene with the second-best DAP-seq peak was added to the analysis. Candidate TF binding motifs in the sets of orthologous upstream regions were identified by the SignalX program using a simple iterative procedure described previously (25). For the TF binding site prediction, *Rhodanobacter* genomes encoding the studied TF were scanned with position-specific weight matrices (PWMs) representing the candidate TF binding motifs using the GenomeExplorer software package (29). The threshold for the site search was defined as the lowest score observed in the training set. Evolutionary conservation of the predicted TF binding sites was calculated as described previously (30).

### Validation of DAP-seq analysis using RT-qPCR

RNA extraction was performed as per the protocol described by (31,32). Briefly, cultured *R. denitrificans* FW104-10B01 wild-type (WT) and FW104-10B01 tn::*clp* strains were harvested at mid-log phase for RNA isolation. Mid-log phase for the two WTs were reached at OD_600_ ~0.15 and for tn::*clp* at OD_600_ ~0.075. To ensure equivalent biomass across samples, cultures were harvested based on a fixed number of OD_600_-units (OD_600_ × volume). Correspondingly, 5 mL and 10 mL cultures were collected for the WT and tn::*clp* strains, respectively, for comparable total cell biomass for RNA extraction. Cultures were centrifuged for 5 min at 4 °C to form pellets that were re-suspended in 0.5 mL of lysis buffer (30 mM Tris, 10 mM EDTA, 10 mg/ml lysozyme, pH 6.2), followed by incubation at 37 °C for 30 min. 1 ml TRIzol (Invitrogen) was added to the lysed bacterial pellets, and incubated at room temperature for 10 minutes. 200 μl of chloroform were added to the lysates, and vortexed before being centrifuged at 15,700 x g for 20 min at 4 °C for phase separation. The aqueous phase (400 μl) of each sample was transferred to new tubes, followed by the addition of 400 μl of isopropanol. The mixtures were incubated at RT for 10 min, followed by centrifugation at 15,700 x g for 20 min at 4 °C. The supernatants were discarded and the pellets were washed with 75% ethanol at 7500 x g for 5 min at 4 °C. The supernatants were discarded, and the pellets were allowed to dry at RT before being re-suspended in 40 μL of RNase-free water and treated with Turbo DNase kit (Thermo Fisher, Waltham, MA, USA) and incubated at 37 °C for 20 min. The qualitative and quantitative analysis of the RNA was performed using Nanodrop 1000 Spectrophotometer (Thermo Fisher, Waltham, MA, USA). RNA extraction was performed for three biological replicates for each strain.

One microgram of total RNA was used as a template for cDNA synthesis using QuantiTect Reverse Transcription kit (Qiagen, Redwood City, CA, USA). The RT-qPCR reaction was set-up in a 20 μl total reaction volume, containing cDNA template, SsoAdvanced Universal SYBR Green Supermix, with specific primers on CFX96 Touch Real-Time PCR system (Bio-Rad, Hercules, CA, USA). The PCR amplification procedure was carried out at 98 °C for 10 minutes, followed by 40 cycles of 98 °C for 10 seconds, 60 °C for 1 minute. The gene expression levels of all samples were normalized using the endogenous control gene (16S rRNA), and fold expression was calculated according to the 2^-ΔΔCt^ method (33). The experiments were performed in three biological replicates. Student’s t-test was employed to compare the means across groups in the RT-qPCR validation experiments, and differences were considered statistically significant at *p* < 0.05. The complete list of selected target genes with their annotated functions and primers is available in **Supplementary Table 3**.

### Data Visualization

Data obtained from the DAP-seq and comparative genomics pipelines were processed for visualization using the Tidyverse (34) suite of packages (version 2.0.0) in R (version 4.5.1). Several additional packages were used to generate panels for the figures of this manuscript. The visual representation of the genomic positions of TFs and their targets in *R. denitrificans* FW104-10B01 and *R. thiooxydans* FW510-R12 was made using the Circlize package (35) (version 0.4.16). The UpSet visualization of the number of targets of each TF across both genomes was made using the UpSetR package (36) (version 1.4.0). The visualization of the regulatory networks was generated using Cytoscape (37) (version 3.10.4) in conjunction with the RCy3 API (38) (version 2.28.1). Functional annotation of the identified target genes was performed using eggNOG-mapper (version emapper-2.1.12) (39) based on eggNOG orthology data (40). Sequence searches in the eggNOG workflow were performed using DIAMOND (41). Aside from these, as mentioned above, the motif logos obtained in this study were generated using the MEME suite (version 5.3.0) (22). Individual figure panels were assembled into the final version of figures present in this manuscript using Inkscape (version 1.4.2) (42). The data used to generate the figures are available in **Supplementary File 2** and **Supplementary File 3**.

## Results & Discussion

### Identification and Analysis of Predicted TF-Target Gene Interactions from DAP-seq

DAP-seq was conducted to identify TF target genes on a genome-wide scale in two *Rhodanobacter* strains. (**Figure 1**).

**Figure 1.**
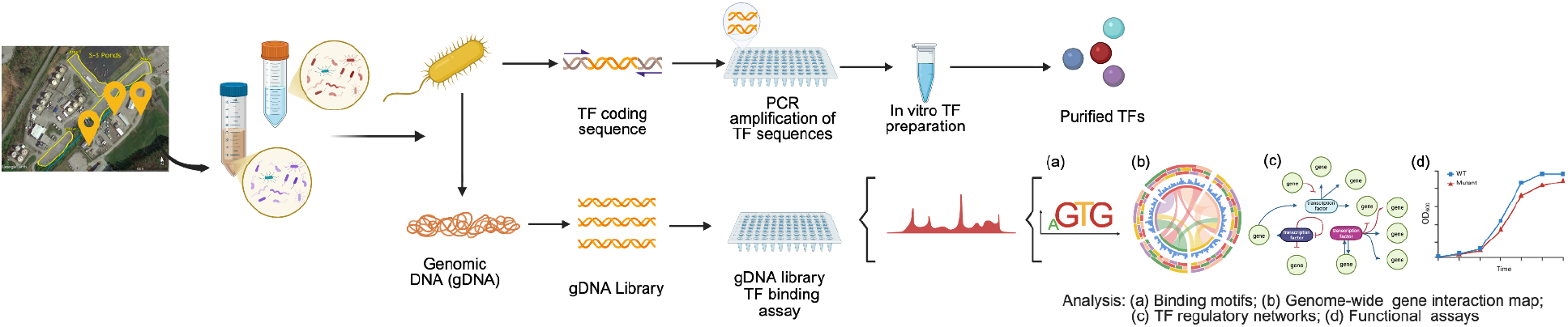
Schematic illustration of the DAP-seq protocol and analysis performed to identify the genome-wide TF binding sites for both *Rhodanobacter* strains. Previous efforts for cultivating microorganisms from the Oakridge Field research site led to the recovery of several isolates. Of these, genomic DNA extracted from two *Rhodanobacter* strains were used in a DAP-seq analysis with a total of 91 transcriptional factors, leading to identification of binding motifs and regulatory networks. Figure generated using Biorender.

DAP-based methods originate from DAP-chip (43) and enable the investigation of TF-DNA interactions under controlled *in vitro* settings, This facilitates studies in organisms for which prior knowledge of regulation is limited. As such, DAP-seq has been used to accelerate gene regulation research in plants, which orchestrate complex transcriptional responses across thousands of TFs in different tissues (44–46) and non-model bacteria, including *Brucella melitensis (47), Clostridium thermocellum* (48), and *Novosphingobium aromaticivorans* (49). In the present study, DAP-seq generated a large dataset of sequencing peaks for several TFs from the two *Rhodanobacter* strains. A total of 91 TFs from 17 TF families were examined for DNA binding, leading to a primary dataset of enriched peaks that comprises over 3,200 sequencing peaks for 30 different TFs in *R. denitrificans* FW104-10B01, and 4,500 for 33 different TFs in *R. thiooxydans* FW510-R12. Despite the substantial output of this approach, the distribution of quality hits per TF was uneven across the dataset. Notably, 16 of the investigated TFs of *R. denitrificans* 10B01, and 12 of *R. thiooxydans* R12, did not have any associated peaks that met our analysis cut-off threshold, and the TFs LRK54_RS11585 of strain 10B01 and LRK53_RS16970 of strain R12, both annotated as Clp, were highly enriched and represented ~10% of the total peaks in both genomic samples.

After curation, we identified 12 TFs from FW104-10B01 and 22 TFs from FW510-R12 that produced satisfactory DAP-seq peaks in upstream regions of protein-coding genes in the primary dataset. Out of these, we excluded three TFs from our subsequent analyses: One TF (LRK53_RS04505), a truncated homolog of the methionine repressor MetJ, generated over 600 DAP-seq peaks in both genomic DNA samples but lacked a discernible sequence-specific binding motif. The other two TFs (LRK54_RS05985 and LRK53_RS17390) were also excluded because they only bound to a single site within a filamentous prophage in the FW510-R12 genome. Therefore, the final subset of the DAP-seq dataset used in this study is composed of 31 TFs, 25 of them belonging to 13 ortholog groups from both *Rhodanobacter* and six TFs unique to either strain, that were selected for further investigation (**Figure 2a**).

**Figure 2.**
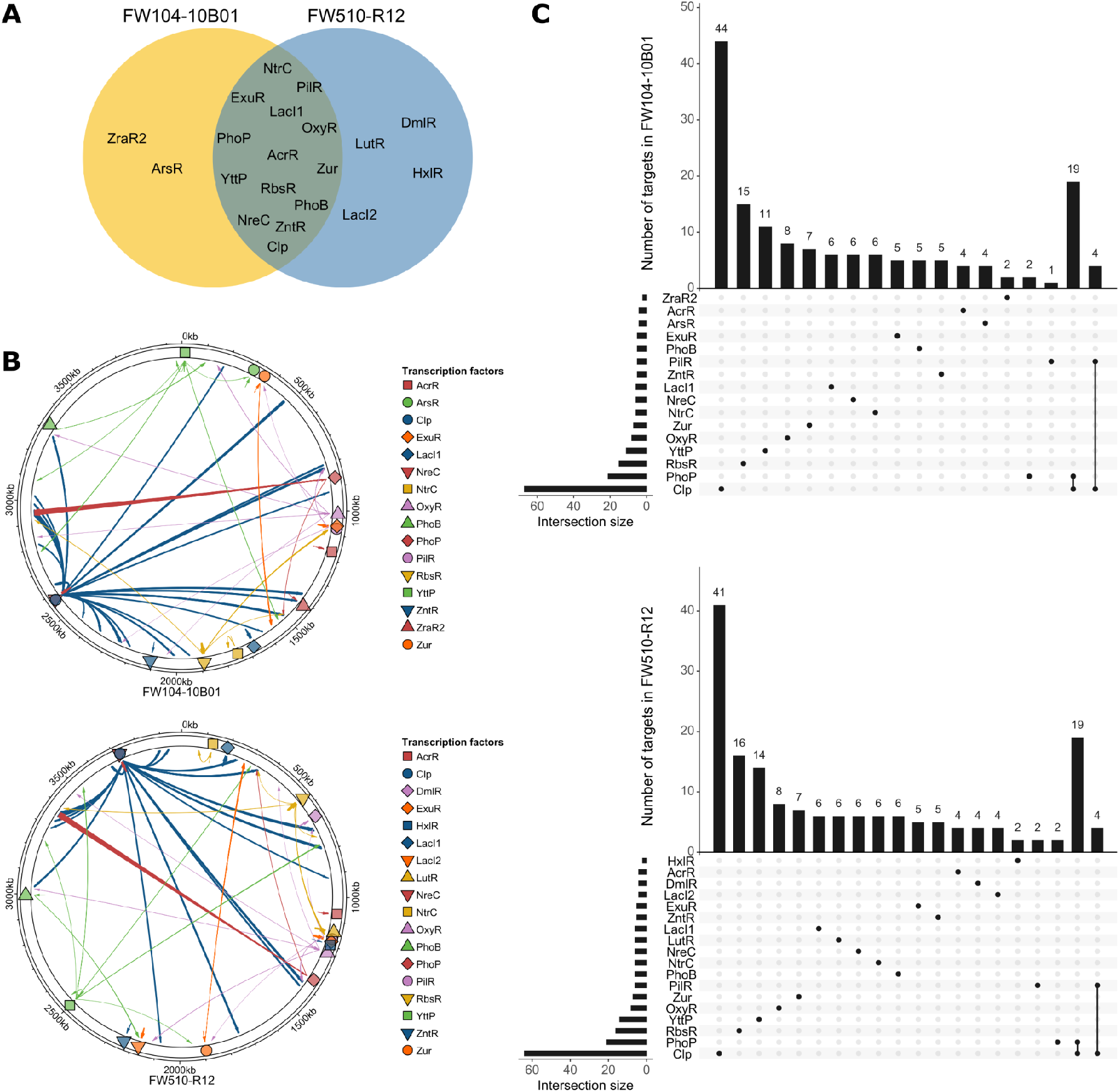
(a) Venn diagram depicting the conserved and strain-specific TFs present in both *Rhodanobacter* strains investigated in this study. (b) Genomic coordinates of the TFs studied (represented by different colored shapes along the genomic ruler) in the chromosome of *R. denitrificans* FW104-10B01 (top panel) and *R. thiooxydans* FW510-R12 (bottom panel). In this visualization, each TF is connected to their genomic targets by an arrow. TFs that auto-regulate or regulate targets in their close genomic proximity are shown as shorter arrows (c) UpSet plots indicating the number of identified target genes for the studied TFs in the genomes of *R. denitrificans* FW104-10B01 (top panel) and *R. thiooxydans* FW510-R12 (bottom panel). In both plots of panel C, the dark circles represent a specific set of one or more TFs. The vertical bars above the circles indicate the number of target genes regulated by a given set, while the horizontal bars to the left depict the number of genes regulated across all sets for a given transcription factor (intersection size). Of note, two TFs belonging to the LacI family (LRK54_RS07715/LRK53_RS00805 and LRK53_RS09640) had no previous associated genomic annotations for the studied organisms, and are referred to as LacI1 and LacI2, respectively, throughout this manuscript.

To obtain a deeper understanding of these TFs, we used a comparative genomics approach to further augment the DAP-seq findings. Comparative genomics is a powerful *in silico* technique (50), and in this work, we implemented it by utilizing the best peaks reported in the DAP-seq dataset for a given TF alongside with genomic data from 82 *Rhodanobacter* spp. to predict binding motifs, and then utilized this motif to further filter the DAP-seq results (**Methods**). While there are a few genes with comparative genomics predictions and no DAP-seq peak, in most cases we found a good correspondence between the two approaches and found that our analysis benefitted from employing both synergistically.

### Predicted TF binding Motifs From DAP-seq and Comparative Genomics

Data from our combined DAP-seq/comparative genomics workflow shows that TFs across both of the investigated strains exhibit similar regulation patterns, and most of the identified target genes regulated by common TFs are orthologs. In fact, aside from the TFs found uniquely in either genome, the connectivity between the TFs and targets across *R. denitrificans* FW104-10B01 and *R. thiooxydans* FW510-R12 (**Figure 2b**) is highly similar. This is despite the relative genomic locations varying in the two genomes, possibly due to *R. thiooxydans* FW510-R12 specifically exhibiting lower collinearity, greater genomic rearrangements, and a significantly altered gene order when compared to other *Rhodanobacter* strains isolated from ORR environments (10). The coincidences between TF/target interactions identified in our dataset may be attributed to the high similarity of protein sequences (~90%) and close phylogenetic proximity of both strains (10). For instance, in both strains, the highest number of genes were predicted to be regulated by a global regulator Clp, while the lowest number of genes were found associated with RRs such as PilR and PhoP. In addition to this, Clp and PilR are shown to share a few targets in both strains (**Figure 2c**), including a few hypothetical proteins and LRK54_RS03880/LRK53_RS06010 which putatively encode for pilin. Despite these similarities, TFs such as AsrR and LacI2 were found to be exclusively present in the genomes of either *R. denitrificans* FW104-10B01 and *R. thiooxydans* FW510-R12, respectively, and interact with a few targets each. With this considered, our data supports the notion that, despite being isolated from sites under different selective pressures, both strains under investigation share several of the same regulatory networks with few prominent differences between them.

Our results (**Figure 3**) also suggest that TFs commonly present in both *Rhodanobacter* genomes have conserved motifs with similar nucleotide arrangement and probability of occurrence, underscoring the genomic similarity across the two *Rhodanobacter* species and suggesting conserved properties for TFs such as Clp, RbsR and YttP. Moreover, we found that the consensus sequences of identified motifs were variable, but generally concordant, for several of the TFs when comparing the motif logos generated from DAP-seq data and from the comparative genomics analyses (**Figure 3, Supplementary Figure 1, Supplementary File 4**). For instance, despite the TF binding motifs varying significantly within the Cyclic AMP receptor (CRP) family, these transcription regulators are characterized by a conserved C-terminal helix-turn-helix (HTH) DNA binding domain and generally recognize a symmetrical binding sequence (5’-TGTGA-N6-TCACA-3’) located near or within the promoter regions (51–53). Both the Clp binding motifs in *Rhodanobacter* identified using DAP-seq generated enriched sequences and comparative genomics shared significant similarities with canonical Clp motifs identified by other studies (54,55), including, in *R. denitrificans* FW104-10B01, the prominent presence of a conserved N-terminal ‘GTG’ that was shown to play an essential role in the formation of the DNA-protein complex by *Xanthomonas campestris* Clp (55) (**Figure 3a**). Likewise, the binding motif for the LacI family TF RbsR was found to be highly similar in both strains both in the DAP-seq and comparative genomics analyses (**Figure 3b**), and characterized by a conserved core sequence, 5’-AAACGTTT-3’, that was also reported to be conserved in *Bifidobacterium* sp, *Serratia* sp. and several gamma- and beta-proteobacteria (56–58). Yet, we have found that the quality of motifs identified from the DAP-seq pipeline is usually dependent on the number of different binding locations in the genome.

**Figure 3.**
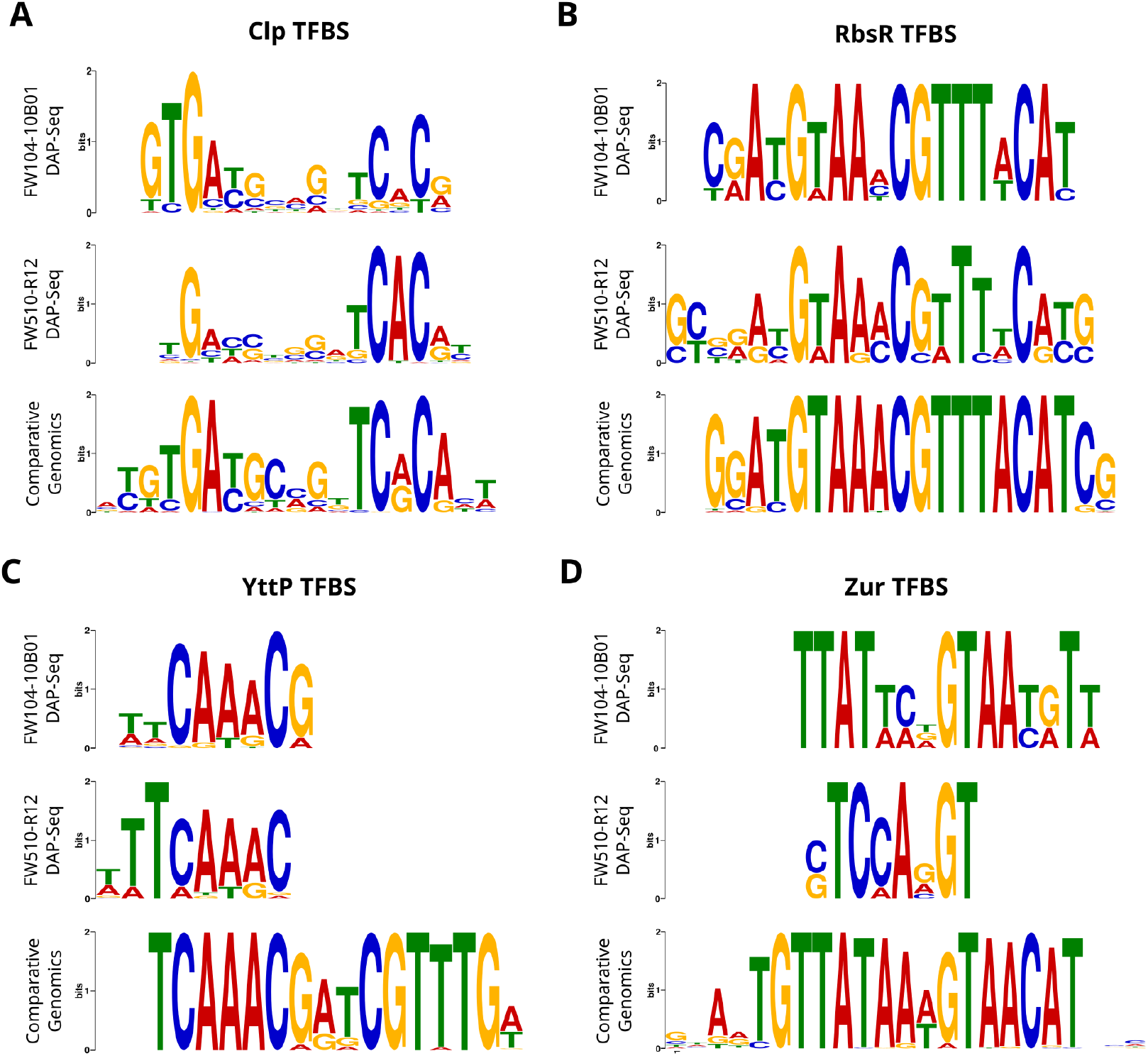
Panels **(a)** – **(d)** depict the motifs identified for four representative TFs using DAP-seq data for both strains (in each panel, top and middle logos) or the broader comparative genomics approach (bottom logos). A full list of inferred motifs is provided in Supplementary Figure 1 and Supplementary File 4.

As seen in **Figure 3c** and **Figure 3d**, the YttP and Zur motifs identified by DAP-seq have low similarity to the respective motifs identified by comparative genomics, which might be caused by the fewer number of binding sites for these TFs in the genomes analyzed by DAP-seq. In fact, for LacI1 and LacI2, no DAP-seq based binding motifs were identified for either *Rhodanobacter*, since these regulators have only one site in both genomes for regulating their targets, which are arranged in operons. Interestingly, while the DAP-seq pipeline identified a peak for PhoB RR upstream of the *pstSCAB-phoU* operon for a putative phosphate transporter in both genomes with predicted a short motif (‘TTTGA’), the comparative genomics approach failed to identify a conserved direct repeat at the summit of this DAP-seq peak. The observed short motif for PhoB in both *Rhodanobacter* genomes is markedly different from the Pho-box of *E. coli*, which consists of a direct repeat of CTGTCAT separated by a 4-bp AT-rich linker (59). PstSCA and PhoU are involved in phosphate regulation and the corresponding operon is a reasonable PhoB target, however the absence of an identified canonical PhoB motif requires additional experimental validation to establish biological relevance of this finding as well as to confirm the binding to the peak identified.

### Functional Analysis of TF-Target Gene Network Clusters

To further characterize the identified TF-target gene regulatory networks in each *Rhodanobacter* strain, we built a visualization linking TFs to their targets and associated gene ontologies utilizing the comparative genomics curated list of targets **(Figure 4**). Our data shows that, for both strains, orthologous TFs regulate similar genes involved in the processes such as uptake and metabolism of essential nutrients, general cellular metabolism, ion metabolism, and biofilm formation. In a similar way, various other conserved TFs regulate genes that also seem to be relevant to the environment found at ORR, including OxyR, which is shown to regulate genes involved with energy conversion and inorganic ion transport and metabolism, and Clp, associated with genes involved in several processes, including cellular motility and the maintenance of the cellular membrane. The CRP family protein Clp functions as global regulatory TF in bacteria and has been reported to be associated with the transcription of almost 300 genes in other organisms such as *Xanthomonas campestris* (60).

**Figure 4.**
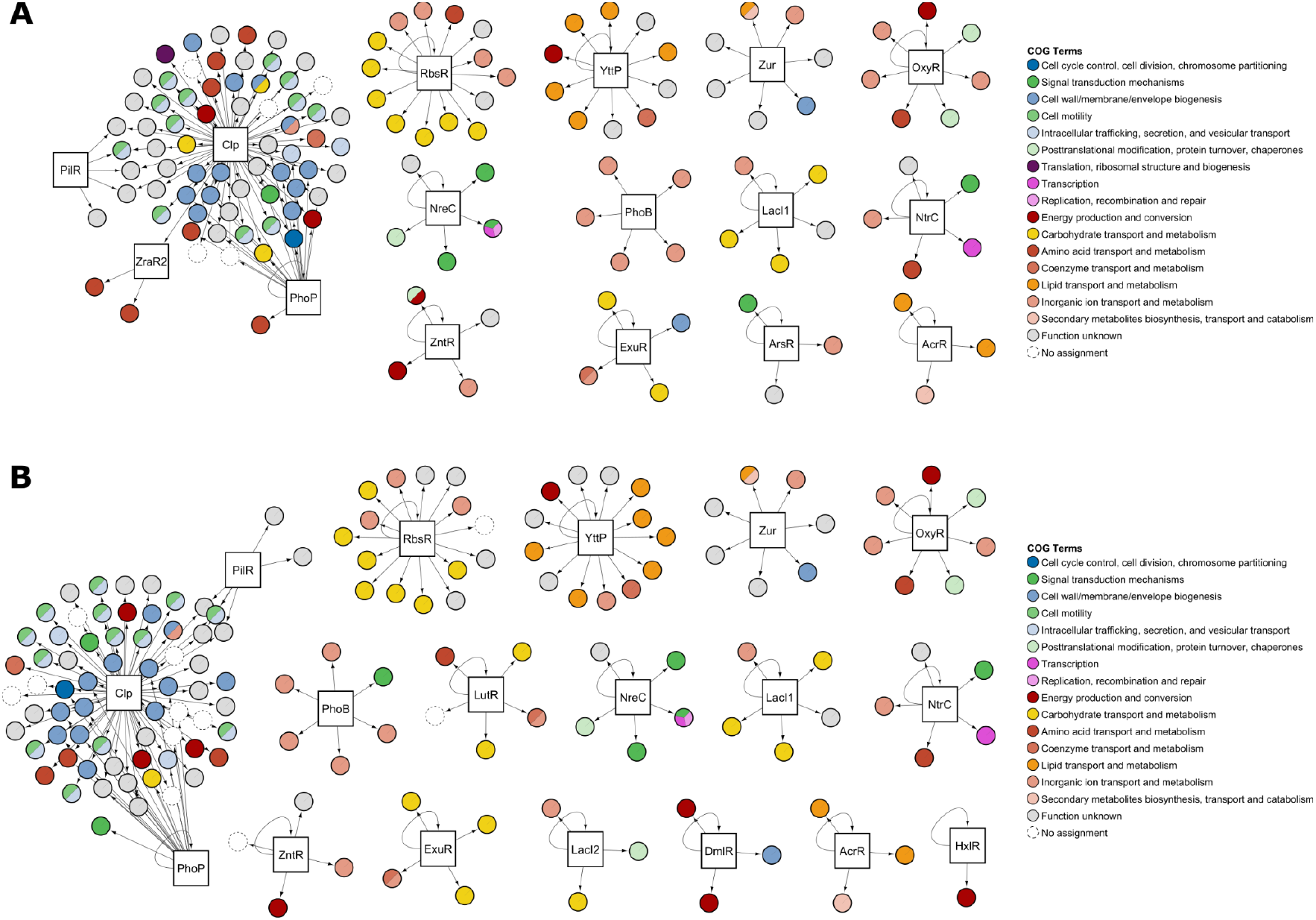
Visualization of *Rhodanobacter* regulatory networks. This figure shows networks of (a) *R. denitrificans* FW104-10B01, and (b) *R. thiooxydans* FW510-R12. In it, each TF investigated in this work is depicted as a central node (white squares), with its identified targets arranged around it (colored circles). Target nodes are color-coded according to their functional categories according to their Cluster of Orthologous Genes (COG) classifications. Targets without associated COG classifications are indicated by white circles with dashed outlines and a “no assignment” label.

In our dataset, Clp was observed to regulate the largest number of genes in both strains, the majority of which related to cellular metabolism, motility, biofilm formation and regulation (**Figure 4**). The ability to form biofilms is a crucial survival strategy for microorganisms in harsh environments (61), and *Rhodanobacter* spp. have previously been shown to use this mechanism as an adaptive response to various stressors such as low pH and metal toxicity, creating microniches with varying pH and/or metal concentrations. Recent research has highlighted that mutations in the *clp* (LRK54_RS11585), *pilR* (LRK54_RS05020) and *fleQ* (LRK54_RS13485) regulatory genes enhance biofilm formation in *Rhodanobacter* under heavy metal stress (17). Our data supports these findings by showing that Clp functions as a key regulator both in *R. denitrificans* FW104-10B01 and in *R. thiooxydans* FW510-R12, controlling genes and response regulators (such as FleQ) involved in flagella and pilin synthesis (**Figure 4a, Figure 4b**), including shared targets with PilR and PhoP, which are regulators associated with pilin and exopolysaccharide formation, respectively.

In contrast to the global scope of ClpTFs such as OxyR were found to be associated with fewer genes closely related in function, which suggest specialized functions. OxyR is a highly conserved transcriptional autoregulator in bacteria that senses peroxide levels and acts as a redox switch, helping to mitigate the harmful effects of reactive oxygen species (ROS) (62). The core OxyR regulon in *E. coli*, primarily includes genes encoding enzymes that directly detoxify ROS like hydrogen peroxide (e.g., catalase, alkyl hydroperoxide reductase), alongside genes involved in iron sequestration, notably *dps*, to mitigate the damaging effects of free iron during oxidative stress (63,64). In *Rhodanobacter*, we observed OxyR to regulate genes such as *ahpC* (alkyl hydroperoxide reductase subunit C) and *ahpF* (alkyl hydroperoxide reductase subunit F), catalase (*cat*), rubrerythrin (*rbr*) and *dps*. The *ahpCF* system in bacteria is the primary and most critical factor responsible for scavenging microscale H_2_O_2_, while *cat* is responsible for degrading large amounts of H_2_O_2_ (65). The gene *rbr*, that codes for the metalloprotein rubrerythrin, has been identified as an alternative to the superoxide dismutase and catalase, and functions by increasing bacterial resistance to H_2_O_2_ and O_2_ (66). It is noteworthy that this gene has been found to play a crucial role in managing oxidative stress in many anaerobic bacteria, and is rarely observed in aerobic bacteria. Given that both *Rhodanobacter* strains are facultative anaerobes capable of switching between different respiratory states in response to fluctuating environmental conditions at the ORR, it can be hypothesized that the presence of *rbr* serves as a survival strategy, helping bacteria to cope with oxidative stress under both aerobic and anaerobic conditions. Together, these proteins contribute to mitigating oxidative stress within the cell under challenging conditions.

Despite the similarities shared between both *Rhodanobacter* strains we observed were key differences in regulation that might point to the specific environments where they were isolated at the ORR site. For instance, *R. denitrificans* strain FW104-10B01 possesses a predicted As(III)-responsive ArsR repressor that has no ortholog in *R. thiooxydans* FW510-R12. ArsR is a transcriptional repressor that binds the promoter of *ars* operon and responds to the presence of arsenite in the environment (67). We found the ArsR of *R. denitrificans* FW104-10B01 to regulate four genes from the *ars* operon, including arsenate (*arsC*) and arsenic (*arsH*) resistance genes. The *arsC* gene encodes a cytoplasmic arsenate reductase that is known to be widespread and horizontally transferred in environments and to have an evolutionarily conserved mechanism of detoxification (68,69, 70). The *arsH* gene encodes a functionally diverse arsenical resistance protein that belongs to the NADPH-dependent FMN oxidoreductase superfamily (71), and it has previously been characterized as a reductase with various substrates, including reduction of azo dyes and some heavy metals including imparting resistance against both arsenite and arsenate (72).

A deeper functional evaluation of these tolerance determinants would better elucidate their roles in enabling survival of these hosts in the original ORR site, the present analysis of the target genes regulated by a set of selected TFs across the two closely related *Rhodanobacter* strains indicated a core regulatory network that is conserved between the two species, and suggested the presence of regulons that are crucial in their adaptation to specific ORR environments. Collectively, our results suggest that the TFs selected in this work primarily regulate genes involved in nutrient uptake, cellular metabolism, motility, and the response to stress, particularly oxidative and inorganic ion stresses.

### Validation of DAP-seq/ Comparative Genomics Predicted TF Regulation Using RT-qPCR Analysis

To validate the *in vivo* function of the most prominent regulator revealed in this study, specifically the global regulator Clp, we examined the effect of its absence on the expression of selected target genes in *R. denitrificans* FW104-10B01. To confirm physiological relevance of this TF, we employed RT-qPCR to measure the expression levels of four putative Clp targets, *pilM, pilA, pilZ*, and *fleQ*, in the WT FW104-10B01 strain and a *clp* transposon mutant (tn::*clp*).

As shown in **Figure 5**, when both WT and the tn::*clp* mutant were grown under normal conditions in R2A media, the relative gene expression of all the four target genes was almost twofold lower in the mutant when compared to the WT, indicating that the absence of Clp alone can affect the expression of these four genes. Notably, the selected target genes are functionally diverse, and while *fleQ* is a transcriptional regulator, the genes *pilM, pilA* and *pilZ* codify proteins that participate in formation and assembly of type IV pilin (T4P) structure, thin and flexible filaments found on the surface of a wide range of Gram-negative bacteria (73). Previous studies have also shown Clp as an essential factor behind T4P-driven motility required for bacterial movement and biofilm formation (54,74), and that Clp directly binds to the promoter sequence upstream of the *pilA* and *pilMNOPQ* operon genes that encode key T4P structural components, leading to the transcriptional activation of these T4P genes, facilitating the formation of T4P and twitching motility of *Lysobacter enzymogenes* OH11 (54,74). Apart from the role of Clp in T4P formation, Clp has also been associated with the regulation of flagella related genes. Sequence analysis of *fleQ* in *Xanthomonas campestris* revealed the presence of a conserved Clp binding domain, indicating that Clp may regulate the expression of *fleQ*, which further acts as a regulator for genes responsible in flagellar synthesis (75). By establishing a relationship between Clp and its target genes in *R. denitrificans* FW104-10B01, this assay is consistent with our initial assessment of the wide scope of the Clp regulon and reinforces the key role that this TF plays in regulating the transcriptional activation of genes involved in pilin formation and flagellar regulation in this organism, both of which are crucial for bacterial motility.

**Figure 5.**
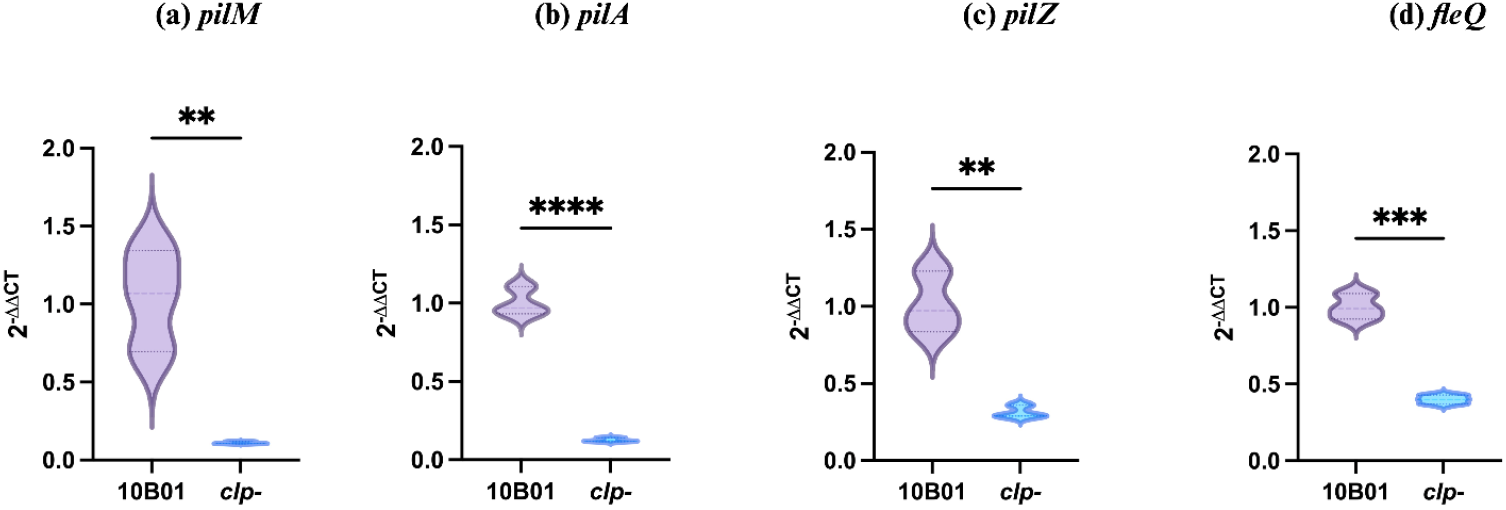
Expression levels of four target genes, namely (a) *pilM*, (b) *pilA*, (c) *pilZ* and (d) *fleQ*, regulated by Clp (a global regulator) as analyzed by RT-qPCR for *R. denitrificans* FW104-10B01 (WT) and the tn::*clp* mutant. For each strain, the relative gene expression of the target genes was normalized to the 16S rRNA housekeeping gene.

## Conclusions

This study the regulatory networks of two closely related *Rhodanobacter* strains by performing DAP-seq and comparative genomics, and analyzed target genes and TF binding motifs for 31 TFs. The resulting dataset revealed remarkably conserved regulatory patterns within the two strains, and the presence of orthologous binding targets for several TFs with important functions in bacterial motility, carbohydrate metabolism, ion transport and metabolism of essential nutrients such as phosphate and nitrogen. Several the targets for specialized TFs were also elucidated. Key differences for targets in commonly present RRs such as OxyR reveal the niche specific regulatory networks for these bacteria. Clp emerged as a key effector of well-structured regulon in this study and we validated its binding using additional assays. Understanding the regulatory landscape in these strains provides a more granular understanding of the signaling systems required for response and survival in the ORR.

Although each DAP-seq peak ideally captures a TF binding event and a potential interaction site for RNA polymerase (48), non-specific binding and experimental limitations, such as missing post-translational modifications in purified TFs, variable binding affinities, or cooperative binding with other TFs, can lead to inaccurate peak calls (76). To mitigate these issues and increase confidence in motif identification, in this study we applied two complementary strategies. First, for each TF with DAP-seq peaks, we generated recurring sequence motifs from two datasets: the full peak sequences and 60-bp regions centered on peak summits. Second, we searched for evolutionarily conserved motifs in upstream regions of 1–2 genes associated with the strongest DAP-seq peaks, as well as in their orthologs across a broad set of *Rhodanobacter* genomes.

DAP-seq is experimental and captures species- and condition-specific binding events, whereas comparative genomics may miss non-conserved or weakly conserved TFBSs. Conversely, DAP-seq may detect non-specific interactions if proper cofactors or protein modifications are absent. For some TFs, the DAP-seq approach alone was not sufficient for the identification of high quality binding motifs. One reason could be a very small number of TFBSs for a protein in the genome, especially for local regulators, thus preventing motif identification. Another issue could also be from non-specific protein binding in the in vitro DAP-seq assay. Thus, for several TFs with weak binding or very few TFBSs (e.g., OxyR, NtrC, NreC, PhoP, MerR, AcrR, LacI1), regulatory predictions relied exclusively on comparative genomics for one or both strains, and additional experimental validation, such as electrophoretic mobility shift and gene expression assays, will be essential to more accurately define their binding sites. The dual approach allowed us to investigate the data from complementary angles and strengthened the robustness of our findings, to enable the identification of TF-target gene interactions in these non-model microorganisms.

## Acknowledgements

The authors thank the Mukhopadhyay group and the ENIGMA team for their constructive feedback regarding this manuscript. We thank Allison Hung (UC Berkeley) careful review of the manuscript and helpful comments. This material is based upon work done in ENIGMA, a Science Focus Area led by Lawrence Berkeley National Laboratory supported by the U.S. Department of Energy, Office of Science, Biological and Environmental Research under Contract Number DE-AC02-05CH11231. The work (https://doi.org/10.46936/10.25585/60008484) was conducted by the US Department of Energy Joint Genome Institute (https://ror.org/04xm1d337), a DOE Office of Science User Facility, supported by the Office of Science of the US Department of Energy under Contract No. DE-AC02-05CH11231.

## Author Contributions

VB, AM developed the project. A. Kakouridis contributed to early conceptualization and sample submissions to the JGI. IB, LB conducted the DAP-seq analysis at JGI. AK conducted comparative genomics. VB conducted experiments and data analysis and initial figures. AC, SP, AK provided experimental support. GMVS analysed data and prepared data figures. VVT, AMD provided reagents and guidance. AM obtained funds and provided supervision. VB wrote the first draft. AK, GMVS and AM helped develop the first draft. All authors contributed to, revised and approved the draft.

