## Supplementary Figure 1 for "Mapping Genome-wide Transcription Factor Binding Sites in two *Rhodanobacter* strains Isolated from Extreme Environments"


### Supplementary Information

### Comparative Genomics Motifs

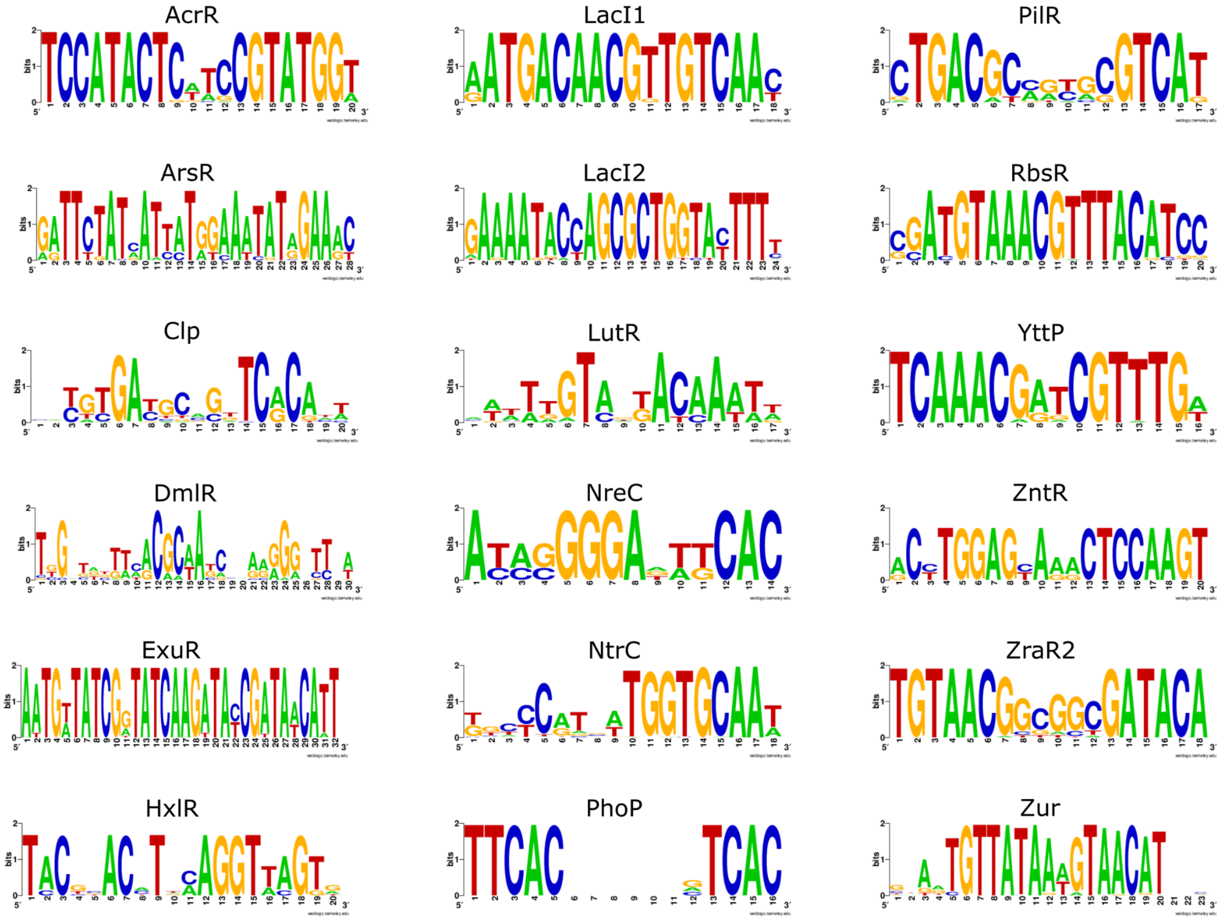


**Supplementary Figure 1.** TF-binding motifs obtained from the analysis of 82 *Rhodanobacter* spp. genomes utilizing the comparative genomics approach.
